# Optogenetics enables real-time spatiotemporal control over spiral wave dynamics in an excitable cardiac system

**DOI:** 10.1101/413658

**Authors:** Rupamanjari Majumder, Iolanda Feola, Alexander S. Teplenin, Antoine A. F. de Vries, Alexander V. Panfilov, Daniël A. Pijnappels

## Abstract

Propagation of non-linear waves is key to the functioning of diverse biological systems. Such waves can organize into spirals, rotating around a core, whose properties determine the overall wave dynamics. Theoretically, manipulation of a spiral wave core should lead to full spatiotemporal control over its dynamics. However, this theory lacks supportive evidence (even at a conceptual level), making it thus a long-standing hypothesis. Here, we propose a new phenomenological concept that involves artificially dragging spiral waves by their cores, to prove the afore-mentioned hypothesis *in silico*, with subsequent *in vitro* validation in optogenetically-modified monolayers of rat atrial cardiomyocytes. We thereby connect previously established, but unrelated concepts of spiral wave attraction, anchoring and unpinning to demonstrate that core manipulation, through controlled displacement of heterogeneities in excitable media, allows forced movement of spiral waves along pre-defined trajectories. Consequently, we impose real-time spatiotemporal control over spiral wave dynamics in a biological system.

## Introduction

Self-organization of macroscopic structures through atomic, molecular or cellular interactions is characteristic of many non-equilibrium systems. Such emergent dynamic ordering often reveals fundamental universalities (***Cross and Hohenberg, 1993***). One example is the occurrence of rotating spiral waves. Spiral waves are found in diverse natural systems: from active galaxies (***Schulman and Seid 1986***), to simple oscillatory chemical reactions ***Belousov (1985)***; ***Zhabotinsky (1991)***, to social waves in colonies of giant honey bees (***Kastberger et al., 2008***), to Min protein gradients in bacterial cell division (***Caspi and Dekker, 2016***), and to the formation of vortices in fluids flowing past obstacles (***Karman, 1937***). While being beneficial to some systems, e.g. slime molds, where they guide morphogenesis, such activity has detrimental consequences for other systems including the heart, where they underlie lethal cardiac arrhythmias (***Davidenko et al., 1990***). Understanding the dynamics of spiral waves in order to establish functional control over a system, has intrigued researchers for many decades. It has been reported that irrespective of the nature of the excitable medium, spiral wave activity organizes around an unexcitable center (core), whose properties determine its overall dynamics (***Krinsky, 1978***; ***Beaumont et al., 1998***). Theorists attribute such particle-like behaviour of a spiral wave to an underlying topological charge, which controls its short-range interaction, annihilation, and the ability to form intricate bound states with other spirals (***Ermakova et al., 1989***; ***Schebesch and Engel, 1999***; ***Steinbock et al., 1992***).

Rotational activity similar to spiral waves can also occur around small structural or functional heterogeneities (e.g. areas of conduction block). In this case, the dynamics of the rotating wave and its spatial position are determined by the location and properties of the heterogeneity. Thus, in theory, by controlling the position and size of spiral wave cores, one can precisely and directly control the dynamics of spiral waves in general. In order to achieve such control, it is therefore logical, to consider as a first step, possible core-targeting via the conversion of a free spiral wave to an anchored rotational activity. To this end, a detailed mechanistic study was performed by (***Steinbock and Müller, 1993***), who demonstrated the possibility to forcibly anchor meandering spiral waves in an excitable light-sensitive Belousov-Zhabotinsky (BZ) reaction system. Furthermore, (***Ke et al., 2015***) demonstrated in a three dimensional BZ reaction setting, that forced anchoring of scroll waves to thin glass rods, followed by subsequent movement of the rods themselves, could enable scroll wave relocation (***Ke et al., 2015***). On a broader perspective, this could have significant meaning for the heart, where controlling the dynamics of scroll waves could add to the treatment of cardiac arrhythmias sustained by such waves.

In cardiac tissue, the analogs of a classical spiral wave and a wave rotating around a heterogeneity, are, respectively, functional and anatomical reentry, both of which are recognized as drivers of arrhythmias. Interestingly, functional and anatomical reentrant waves are closely related to each other. Seminal findings by (***Davidenko et al., 1991***) demonstrated that a drifting spiral wave could anchor to an obstacle and thereby make a transition from functional to anatomical reentry. Conversely, (***Ripplinger et al., 2006***) showed that small electric shocks could unpin a reentrant wave rotating around an obstacle, bringing about the reverse transition from anatomical to functional reentry. Independently, (***Nakouzi et al., 2016***) and (***Zykov et al., 2010***) demonstrated that the transitions between anchored and free spiral states may be accompanied by hysteresis near the heterogeneities. Furthermore, (***Defauw et al., 2014***) showed that small-sized anatomical heterogeneities could attract spiral waves from a close distance, and even lead to their termination if located near an inexcitable boundary. However, to date, all studies dedicated to spiral wave attraction and anchoring involved the presence of anatomically predefined, permanent heterogeneities, or continuous-in-time processes, thereby making it impossible to manipulate spiral wave cores in a flexible, systematic and dynamical manner. In the present study, we propose a new phenomenological concept to demonstrate real-time spatiotemporal control over spiral wave dynamics through discrete, systematic, manipulation of spiral wave cores in a spatially extended biological medium, i.e. cardiac tissue. We establish such control through optogenetics (***Boyden et al., 2005***; ***Bi et al., 2006***; ***Deisseroth, 2015***; ***McNamara et al., 2016***), which allows the creation of spatially and temporally predefined heterogeneities at superb resolution at any location within an excitable medium. Previous studies e.g, by (***Arrenberg et al., 2010***; ***Bruegmann et al., 2010***; ***Jia et al., 2011***; ***Bingen et al., 2014***; ***Entcheva and Bub, 2016***) and (***Burton et al., 2015***), demonstrate the power of optogenetics in cardiac systems. Thus, the same technology was chosen to strategically exploit fundamental dynamical properties of spiral waves, like attraction, anchoring and unpinning, to discretely and effectively steer spiral wave cores along any desired path within an excitable monolayer of cardiac cells. These findings are highly relevant for understanding non-linear wave dynamics and pattern formation in excitable biological media, as they enable, for the first time, real-time discrete dynamic control over processes that are associated with self-sustained spiraling phenomena, e.g. reentrant electrical activity, cAMP cycles and movement of cytosolic free Ca2+, to name a few. In particular, in the heart, tight control of spiral waves may allow restoration of normal wave propagation.

## Results

Self-sustained spiral waves can be actively generated in most natural excitable media. In this study, we induced spiral waves of period 60 ± 5 ms *in silico* and of period 63 ± 11 ms *in vitro* in confluent monolayers of optogenetically modified neonatal rat atrial cardiomyocytes (see Methods for details). Targeted application of light to the monolayers led to the sequential occurrence of two events: (i) creation of a spatially predefined temporal heterogeneity (i.e. a reversible conduction block) near the core of a spiral wave, and (ii) emergence of a wave from the spot of illumination. In a previous study (***Feola et al., 2017***), we demonstrated the possibility to create such a temporal heterogeneity with optical control over the size, location and duration of the block. Here, we show how creation of a light-induced block close to the core of a spiral wave, can attract the spiral wave tip, causing it to eventually anchor to the block. Subsequent movement of the light spot to different locations within the monolayer, results in dragging of the spiral wave along a predefined pathway of illumination, thereby giving rise to, what we call Attract-Anchor-Drag-based (AAD) control. Figure 1A illustrates a schematic diagram of the experimental set up that we used for our *in vitro* studies, whereas, the sequence of light patterns used for real-time AAD control is described in Figure 1B. Figures. 1(C1-4) and 1(D1-4) show representative examples of AAD control of spiral wave dynamics *in silico* and *in vitro*, respectively, where the spiral wave core is dragged along a predefined 9-step triangular path (see also Video 1). Our results demonstrate the possibility to capture a spiral wave, to manipulate its trajectory in a spatiotemporally precise manner (Figures. 1 C2-3 and D2-3), and, finally, to release it in order to rotate freely again (Figure 1 C4, D4). Thus, we prove that it is indeed possible to directly and precisely manipulate the spatial location of a stable spiral wave core, and thereby overcome the constraints imposed by internal and external factors that shape the natural meander and drift trajectories. Such control is very important because it provides a direct and effective handle over the spatiotemporal patterning of the system, which would affect its general functionality. Natural systems currently suffer from the lack of such a handle (***Mikhailov and Showalter, 2006***; ***Mikhailov and Loskutov, 2013***). Spiral wave dragging seeks to fill this lacuna at the simplest level.

**Figure 1.**
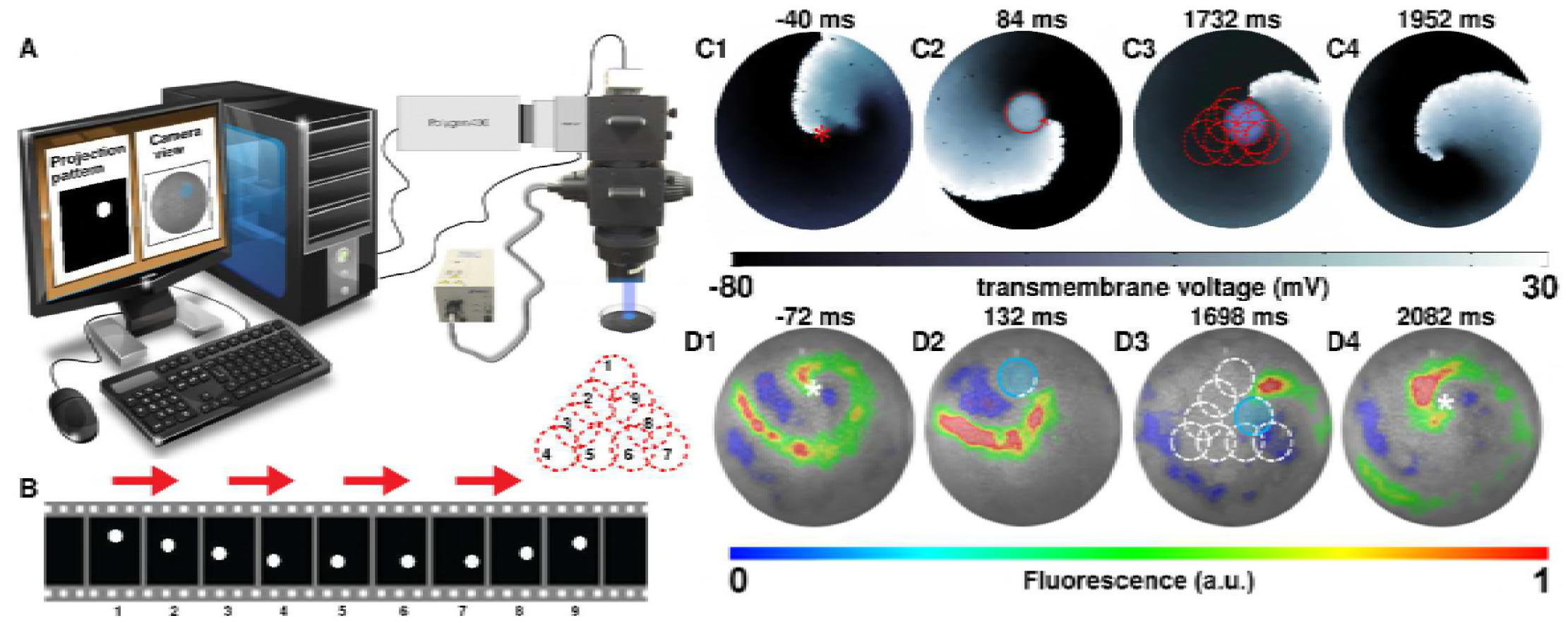
Attract-Anchor-Drag (AAD) control of a spiral wave core along a triangular trajectory. (A) Schematic diagram of our *in vitro* set up, showing how we project light patterns on an optogenetically modified monolayer. (B) The sequence of light spots that constitute the desired triangular trajectory of the spiral core. (C1-C4) *In silico* AAD control data recorded at subsequent times; (D1-D4) *In vitro* corroboration of our *in silico* findings (n=9). In panels (C) and (D), the current location of the applied light spot is indicated with a filled blue circle, whereas, the schematic movement of the tip of the spiral wave, as it is anchored to the location of the light spot at previous time points, is indicated in each frame by means of dashed red (*in silico*) and white (*in vitro*) lines. The location of the phase singularity of the spiral wave is marked in the first and last frame of panels (C) and (D) with a red (*in silico*) or white (*in vitro*) asterisk. a.u., arbitrary units. A video demonstrating the process of dragging a spiral wave core along a triangular trajectory is presented in Video 1.

An immediate application of such a process, that follows dynamic manipulation of spiral waves, involves using this technique to terminate complex spiral wave activity. We first focus on removing a single spiral wave *in silico* (Figure 2 A1-5; see also Video 2). With a 5-step drag sequence of circular light spots of 0.275 cm diameter and 250 ms illumination time per spot, we demonstrate the possibility to remove a spiral wave from a monolayer, by capturing its core somewhere near the middle of the simulation domain and subsequently dragging it all the way to the inexcitable boundary on the left, causing the phase singularity to collide with the border and annihilate. The representative trajectory of the spiral wave, as it is dragged to termination, is shown in Figure 2 (A2-4) with dashed red lines. The direction of movement of the spiral is indicated in each frame with red arrows. Inspired by the outcome of our *in silico* experiments, we tested the same principle *in vitro*, with a 5-step drag sequence of circular light spots with similar characteristics as those applied *in silico*. Our *in vitro* findings corroborate the *in silico* results (Figure 2 B1-5; see also Video 2). The representative trajectory of this spiral wave, as it is dragged to termination, is shown in Figure 2 (B2-4) with dashed white lines. In each frame of Figure 2, the location of the light spot is marked with a transparent blue circle.

**Figure 2.**
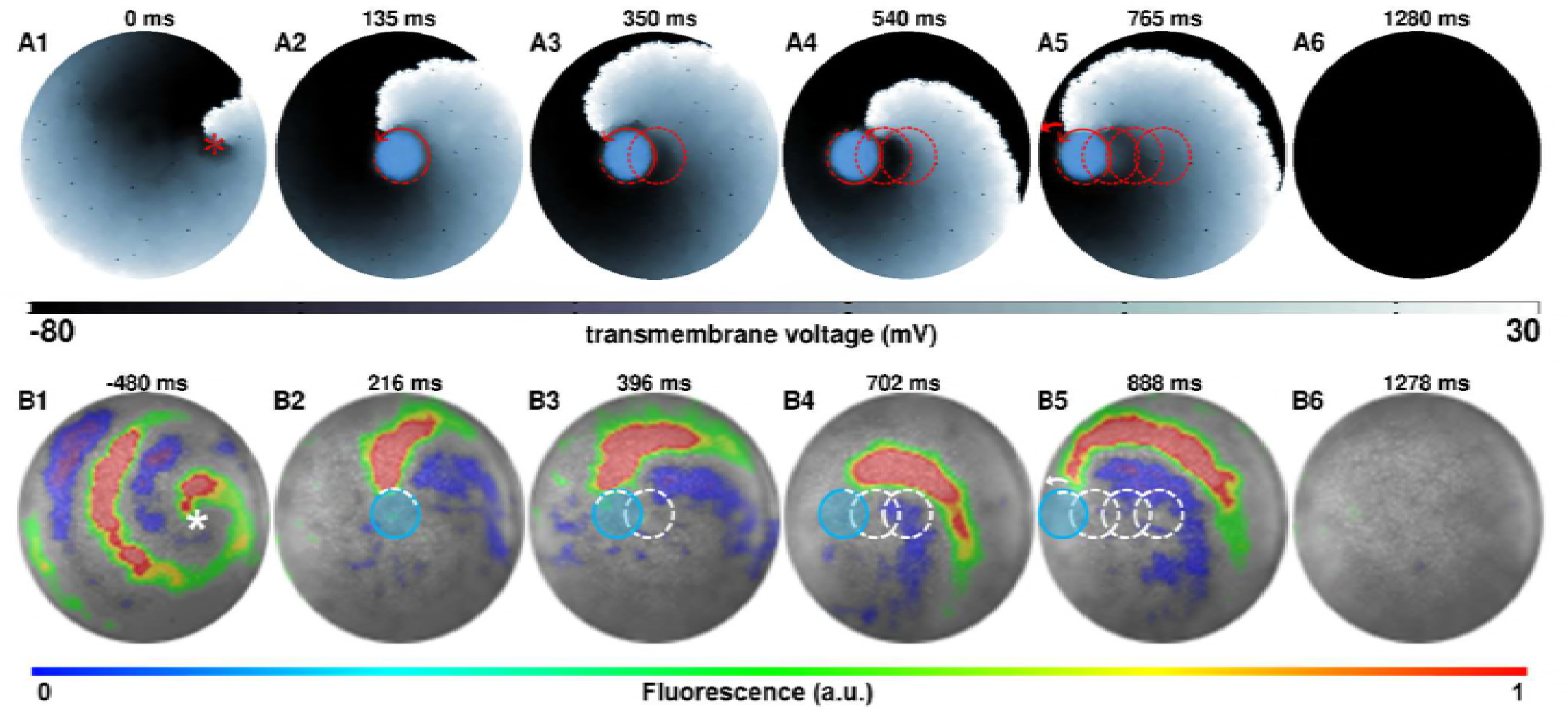
AAD control of a spiral wave core in favor of termination. The upper panel (A1-A6) shows successful removal of a spiral wave *in silico*, by capturing its core from the center of the simulation domain, and dragging it to the left boundary in a stepwise fashion (*τ*_*light*_ = 250 ms and *τ*_*dark*_ = 1 ms per spot, period of the reentry around the spot is 78 ± 2 ms). The lower panel (B1-B6) shows *in vitro* proof of the results presented in panel (A) (n=9). For each light spot, the current location of the applied light spot is indicated with a filled blue circle (*τ*_*light*_ = 250 ms and *τ*_*dark*_ = 15 ms, period of the reentry around the spot is 90 ± 12 ms). The schematic movement of the tip of the spiral wave, as it is anchored to the location of the light spot at previous time points, is indicated in each frame by means of dashed red (*in silico*) and white (*in vitro*) lines. The location of the phase singularity of the spiral wave is marked in the first frame of each panel with a red (*in silico*) or white (*in vitro*) asterisk. a.u., arbitrary units. A video demonstrating the complete process of dragging a spiral wave core from the center of the monolayer to the left border, causing its termination, is presented in Video 2. (Time t=0 ms denotes the moment when the light is applied.).

Having proven the possibility to drag single spiral waves to inexcitable tissue borders in favor of termination, we attempt to develop an in-depth *in silica* qualitative understanding of the parameters that play a crucial role in the dragging process. We focus on the previously described case: linear 5-step dragging from the center of the simulation domain to a point on the periphery, with continuous illumination, and start by investigating the dependence of the probability of successful dragging (*P*_*drag*_), on the diameter (*d*) of a continuously illuminated moving spot (i.e. without a finite time gap between successive applications of light). Our study reveals a positive correlation between *P*_*drag*_ and *d*. At *d ≲* 3 mm, the minimum time (*τ*_*min*_) required for relocation of a spiral wave to the next position decreases abruptly with increasing *d*. However, at *d >* 3 mm, *τ*_*min*_ decreases slowly, trending towards a possible saturation value. These results are illustrated in Figure 3. Traces of the spiral tip trajectory (indicated by means of red lines on top of a representative frame depicted in Figure 3 A1-A3) demonstrate that the ease with which a spiral core can be dragged, from the center of the domain to a border, increases with increasing *d*. At small *d*, the spiral tip is dragged along a cycloidal path, in which, the diameter of the loop is 𝒪(*d*). As *d* increases, the spiral tends to translate linearly, without executing rotational movement about the core.

**Figure 3.**
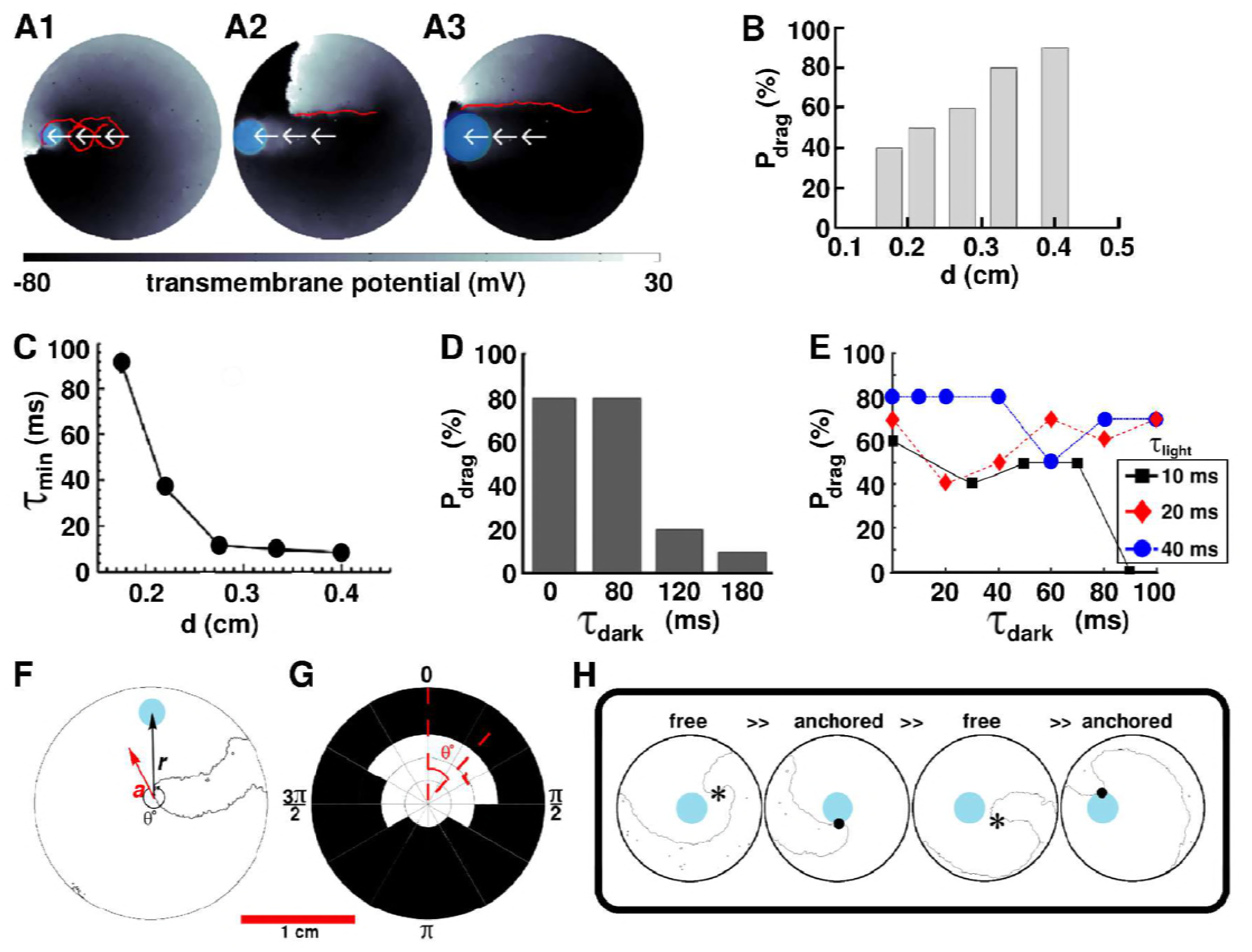
AAD control by continuous illumination of circular light spots of different sizes and durations. Panel A shows representative dragging events as a spiral wave core is relocated from the center of the simulation domain to the periphery, with light spots of diameter *d*=0.175, 0.275 and 0.4 cm, respectively. In each case the real trajectories of the spiral tip is marked with solid red lines on top of the voltage map. When *d* is large, the spiral tip exhibits a tendency to move along a linear path, as opposed to the cycloidal trajectory at small *d*. With *d* = 0.275cm, (B) probability of successful dragging (*P*_*drag*_) increases, and (C) spiral core relocation time (*τ*_*min*_) decreases, with increasing *d*. In each of the afore-mentioned cases, the dragging process is illustrated with zero dark interval (*τ*_*dark*_.) and minimal light interval (*r*_*light*_). (D) With finite non-zero *τ*_*dark*_, *P*_*drag*_ increases with shortening of *τ*_*dark*_, for light stimulation of a fixed cycle length (200 ms). (E) Dependence of *P*_*drag*_ on *τ*_*dark*_, for 3 different light intervals (*r*_*light*_). (F) Schematic representation of the drag angle *θ* and the distance (*r*) between the starting location of the spiral tip and the location of the applied light spot. (G) Distribution of *P*_*drag*_ at different *θ* and *r*, thresholded at 50%. (H) AAD control is effectuated by alternate transitions between functional (free wave) and anatomical (anchored wave) reentry. The phase singularity of the free spiral is marked with a black asterisk, whereas, the point of attachment of the anchored wave to the heterogeneity is marked with a black dot.

To investigate the conditions that allow spiral wave dragging with discrete light pulses, we apply a series of circular light spots of duration *τ*_*light*_. These spots appear sequentially in time, and are physically separated in space, with gradually decreasing distance from the boundary. For simplicity, we apply all spots along the same line, drawn from the center of the simulation domain to the boundary. We define the time gap between successive light pulses (when the light is off everywhere) as the dark interval *τ*_*dark*_. Thus, effectively, our optical stimulation protocol is periodic in time (with period *τ*_*light*_ + *τ*_*dark*_.), but not in space. We perform two sets of studies for the case of linear dragging. In the first set, we fix *τ*_*light*_ + *τ*_*dark*_ at 200 ms and tune *τ*_*dark*_. In the second set, we fix *τ*_*light*_ (3 different values) and tune *τ*_*dark*_. Our results indeed demonstrate the occurrence of AAD control with discrete illumination (*τ*_*dark*_ ≠ 0 ms). Figure 3 D shows, for example, that at *τ*_*dark*_ = 0, 80, 120, and 180 ms, respectively, *P*_*drag*_ decreases from 80% to 20% to 10%. Interestingly, *τ*_*dark*_ = 80ms appears to be as effective as continuous illumination (*τ*_*dark*_ = 0 ms, Figure 3 D). Thus, our findings confirm that, for a fixed cycle length of optical stimulation by a spot of light of diameter *d* = 0.275 cm, *P*_*drag*_ increases with decreasing *τ*_*dark*_

If, however, we tune to explore the dependence of *P*_*drag*_ on *τ*_*ight*_ at flexible cycle lengths, there is no clear relationship between *P*_*drag*_ and *τ*_*light*_ (Figure 3E), despite an increase in the ease of attraction of the spiral tip towards the illuminated spot at short *τ*_*dark*_. In general, *τ*_*light*_ *≃* 𝒪(0.1*τ*_*dark*_) does not lead to effective AAD control, whereas, *τ*_*light*_ *≃* 𝒪(10*τ*_*dark*_, *τ*_*dark*_ ≠ 0) does. However, exceptional cases also exist. For example, (i) *P*_*drag*_ can be large (i.e. 70%) when a short light pulse (*τ*_*light*_ = 20 ms) is followed by a long dark interval (*τ*_*dark*_ = 100 ms); and, (ii) combinations of abbreviated *τ*_*light*_ and *τ*_*dark*_ (i.e. *τ*_*light*_ = 20 ms, *τ*_*dark*_ = 20 ms) can lead to lower *Pdrag* than pulse combinations with shorter or longer *τ*_*dark*_.

We find that probability of attraction also depends on the phase of the spiral wave rotation at the moment of light exposure. This dependence, in simple terms, can be quantified as dependence on the angle between the vector along the instantaneous direction of motion of the spiral tip 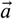 and the vector 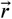 of the displacement of the center of the circular light spot, in polar coordinate system. This is illustrated in Figure 3F. Our results demonstrate that spiral wave dragging occurs most efficiently if the spot of light is applied sufficiently close to the location of the spiral tip and within an angular spread of Δ*θ*_*X*_ in the direction of drift of the spiral core (Figure 3G). Here, *X* denotes the cutoff probability for the occurrence of dragging, i.e. Δ *θ*_50_% refers to an angular spread of Δ*θ* for which *P*_*drag*_ ≥ 50%. A spiral wave cannot be dragged when *r* > 5.63 mm. When 4.7 mm < *r* < 5.63 mm, Δ*θ* = 150°. *P*_*drag*_ increases to 240^°^ when 3.44 mm < *r* < 4.7 mm, and to 360^°^ when *r* < 3.44 mm. Furthermore, to develop a detailed understanding of the physical mechanisms involved in spiral wave dragging, the nature of the process itself is analyzed. Our *in silico* findings indicate that successful control necessitates the spiral wave to make alternate transitions between functional (free) and anatomical (anchored) types of reentry. When a spot of light is applied reasonably close to the core of a spiral wave, within the allowed Δ*θ*, it creates a region of light-induced depolarization that attracts and anchors the core of the free spiral, thereby effectuating a transition from functional to anatomical reentry. When the light spot is now moved to a different location, still within the basin of attraction of the first light spot, the previously depolarized region recovers, forcing the anchored spiral to unpin and make a reverse transition to functional reentry. The larger the value of *τ*_*dark*_, the more visible the transition. Finally, at the new location of the light spot, a zone of depolarization is created, which either attracts and anchors the spiral tip, or produces a new wave of excitation that replaces the existing spiral wave with a new one, still followed by attraction and anchoring. A sequence of *in silico* voltage maps in support of the proposed mechanism, is shown in Figure 3H.

With a detailed understanding of the parameters involved in the process of spiral wave dragging, we explore the possibility to apply this phenomenon to tackle more challenging problems, such as, termination of complex patterns of reentry. We specifically target two patterns: (i) figureof-eight type reentry, and (ii) reentry characterized by multiple spiral waves, i.e. multiple phase singularities. *In silico*, we find that figure-of-eight type reentry can be removed efficiently by dragging the cores of both spiral waves towards each other, till their phase singularities collide, leading to self-annihilation. Figure 4(A1-6) (see also Video 3) show subsequent steps in the process of removal of a figure-of-eight type reentry via AAD control, when the reentry pattern comprises two spirals of opposite chirality, rotating in phase with each other. Figure 4 (B1-6) (see also Video 3) show successful experimental (in vitro) validation of our in silico findings, following the same termination protocol. When the figure-of-eight type reentry comprises two spirals of opposite chirality that do not rotate exactly in phase, the strategy should be to capture the cores of the pair of spirals with light spots of unequal sizes, so as to compensate for the difference in phases, at the very first step of the control method. A representative example of such a scenario is presented in the Figure 4-figure supplement 1.

**Figure 4.**
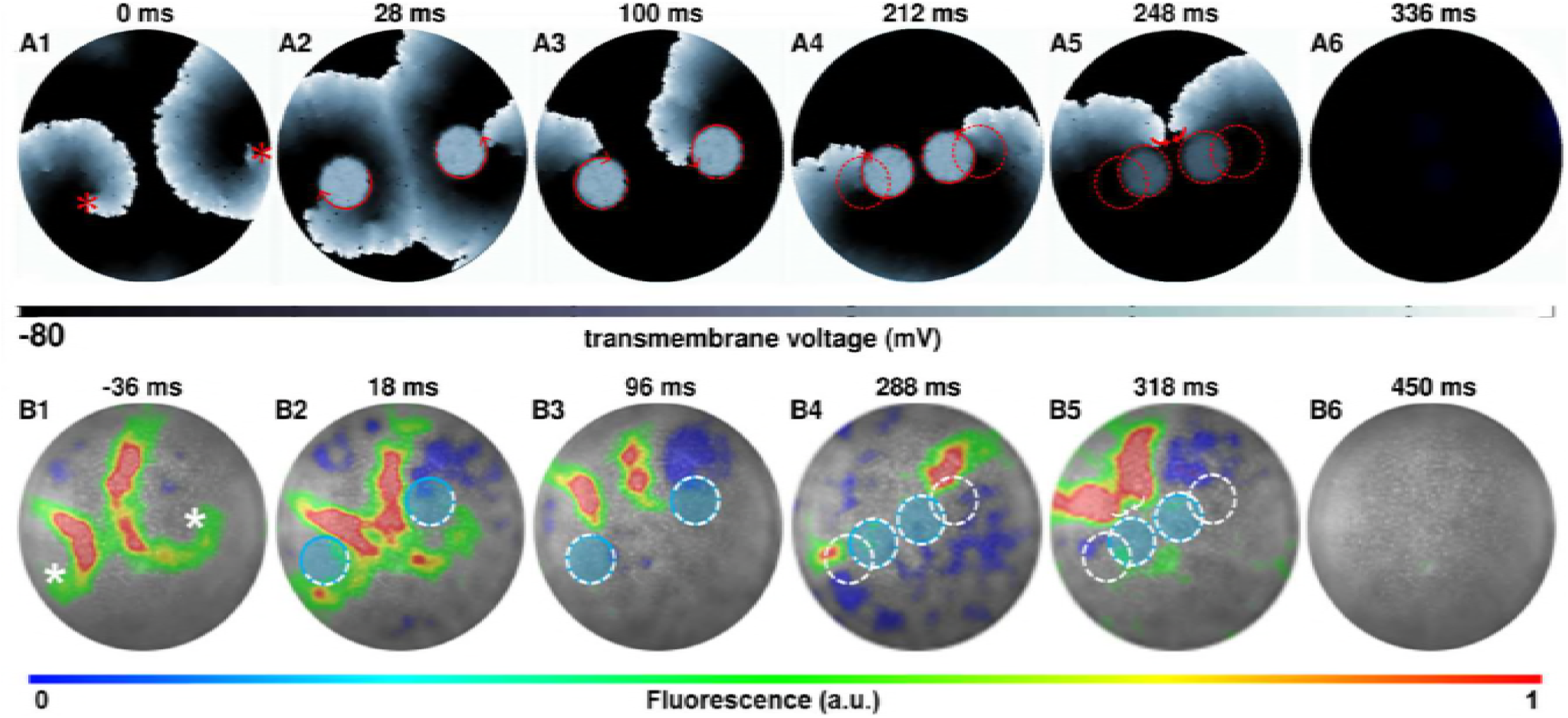
AAD control of a pair of spiral wave cores in favor of termination of figure-of-eight type reentry. The upper panel (A1-A6) shows representative *in silico* voltage maps of the drag-to-termination process, at subsequent times. The lower panel (B1-B6) provides experimental (*in vitro*) validation of the findings presented in panel (A) (n=6). In each case, the locations of the phase singularities are marked on the first frame of each panel with a red (*in silico)* or a white (*in vitro*) asterisk. The current location of the applied spot of light is indicated with a filled blue circle. The schematic movement of the tip of the spiral wave, as it is anchored to the location of the light spot at previous time points, is indicated in each frame by means of dashed red (*in silico*) and white (*in vitro*) lines. The direction of dragging is indicated in panel (A) with red arrows. In our studies, we terminated figure-of-eight type reentry by dragging 2 spiral cores towards each other to make them collide and annihilate. a.u., arbitrary units. A video demonstrating AAD control (with eventual termination) of a figure-of-eight type reentry is presented in Video 3. (Time t=0 ms denotes the moment when the light is applied.)

In order to terminate reentry characterized by multiple phase singularities, we try different approaches. Figure 5 (A1-7) and (B1-7) (see also Video 4) illustrate two examples of such attempts *in silico* with reentry characterized by 3 and 7 phase singularities, respectively. With 3 spiral cores, we follow a strategy which is a combination of the cases presented in Figures 4 and 2, i.e. we capture all 3 cores (Figure 5 A2), drag them towards each other (Figure 5 A3), reduce complexity via annihilation of two colliding phase singularities (Figure 5 A4-5), and drag the remaining spiral core to termination via collision of its phase singularity with the inexcitable boundary (Figure 5 A6-7). However, with higher complexity, there is no unique optimal approach that leads to the termination of reentry. In the example presented in Figure 5 (B1-7), we rely on reducing the complexity of the reentrant pattern itself, via strategic application of 2 light spots, prior to the actual dragging process. New waves emerging from the locations of the applied light spots interact with the existing electrical activity in the monolayer to produce new phase singularities, which collide with some of the preexisting ones to lower the complexity of the reentrant pattern. With reduced complexity, we drag the spiral cores towards each other, till they merge to form a single two-armed spiral, anchored to the applied light spot. We then drag this spiral to the boundary of the monolayer causing its termination. A similar strategy applied *in vitro* enabled successful termination of a complex reentry pattern characterized by 4 phase singularities. This is illustrated in Figure 5 (C1-7). At the instant when the first set of light spots is applied, the reentry is characterized by 4 phase singularities (Figure 5 C1). Note that, of the 4 phase singularities, 2 coincide with the locations of 2 of the applied light spots and 2 are located away from the third light spot (Figure 5 C2). These first 2 phase singularities anchor immediately to the applied light spots. However, the other 2 interact with the new wave generated by the third applied light spot and reduce to 1 phase singularity, which remains free (unanchored) (Figure 5 C3). When the next set of light spots are applied (in a complex pattern of 3 overlapping circles), this phase singularity, displays attraction to the new light spot, thus effectively producing a 3-armed spiral (Figure 5 C4). Finally, when the complex light pattern is replaced by a smaller circular spot of light, the 3-armed spiral loses 2 of its arms through interaction with the emergent new wave from the location of the applied light spot (Figure 5 C5). The single spiral can then be dragged to termination (Figure 5 C6-7) as in Figure 2.

**Figure 5.**
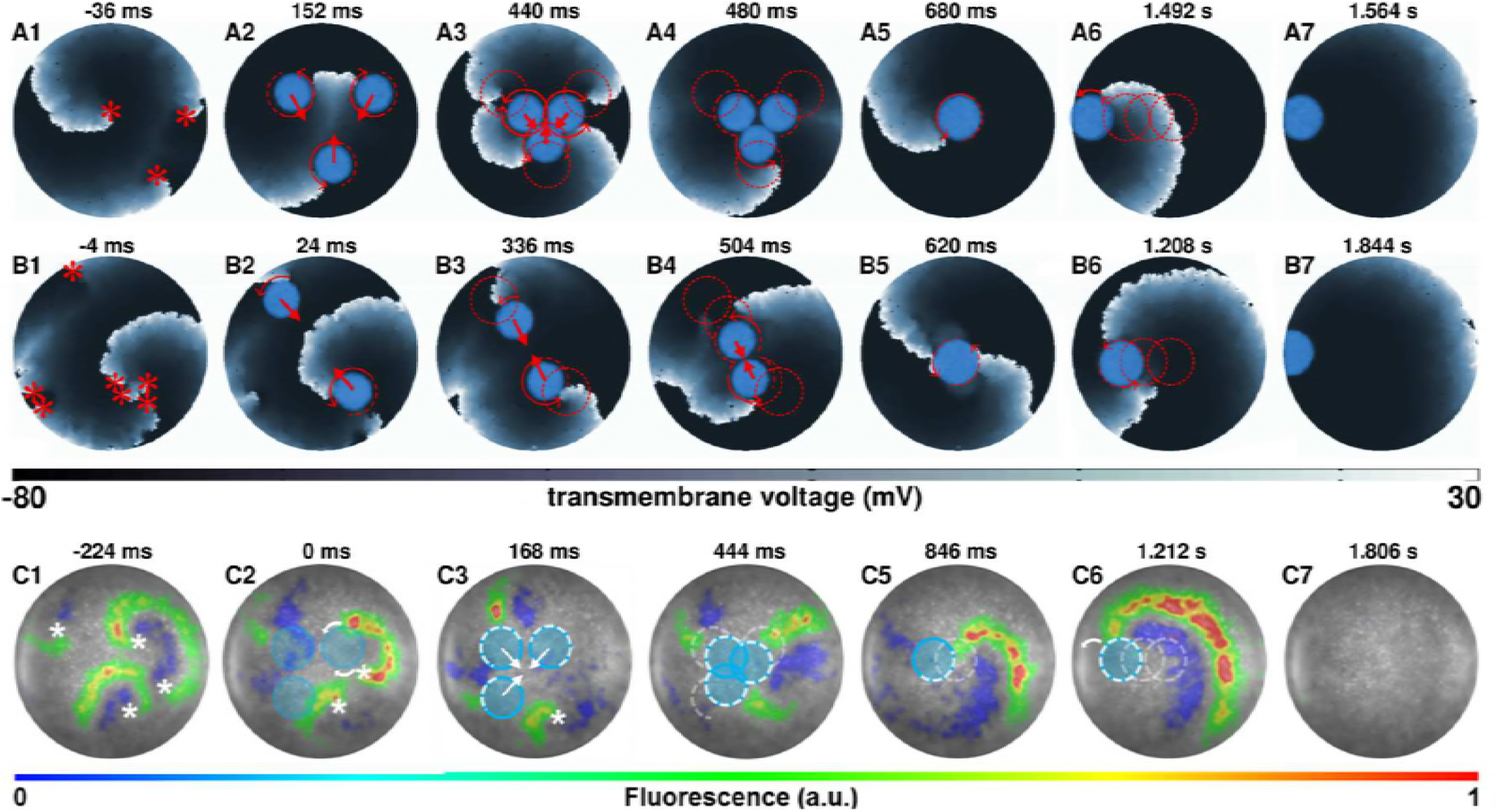
AAD control of multiple spiral wave cores in favor of termination of complex reentrant patterns. Panels (A) and (B) show representative *in silico* voltage maps of the drag-to-termination process for stable reentrant activity with 3 and 7 phase singularities, respectively. Panel (C) shows representative *in vitro* voltage maps of the drag-to-termination process for stable reentrant activity with 4 phase singularities. Number and the position of the light spots, indicated with blue filled circles, are strategically applied to first capture the existing spiral waves, and then drag them to termination, following the same principle as in Figure 2 (A5-6) and (B5-6). In the process it is possible to reduce the complexity of the reentrant pattern through annihilation of some of the existing phase singularities by new waves emerging from the location of the applied light spot. The schematic movement of the tip of the spiral wave, as it is anchored to the location of the light spot at previous time points, is indicated by means of dashed red (white) lines *in silico* (*in vitro*), and the direction of dragging is illustrated with red (white) arrows. Video 4 shows a video of computer simulations demonstrating AAD control of complex reentrant patterns with 3 and 4 spiral waves, in favor of their eventual termination. Video 5 shows the same process, *in vitro*, but for stable reentry with 4 phase singularities. (Time t=0 ms denotes the moment when the light is applied.).

Taken together, our results show that core positions of spiral waves can be strategically manipulated with the help of optogenetics to establish direct spatiotemporal control over a highly nonlinear system. We demonstrate that application of well-timed and spaced light pulses can result in successful dragging of a spiral wave from one location to another within a monolayer, along any desired path and even to cause its termination. Spiral wave dragging can also be employed to terminate complex reentrant patterns characterized by multiple phase singularities, through appropriate AAD control. However, the probability of its occurrence is subject to 2 key parameters: (i) size *d* the tip of the spiral wave, and (ii) drag angle (Δ*θ*)

## Discussion

Spiral waves occur in diverse natural excitable systems, where they impact the functioning of the system in a beneficial or detrimental manner. Given their dynamic and unpredictable nature, direct control over these nonlinear waves remains a longstanding scientific challenge. While it is an established theory that the properties of the core of a spiral wave determine its overall dynamics (***Krinsky, 1978***; ***Beaumont et al., 1998***), the corollary that manipulation of spiral wave cores should lead to full spatiotemporal control over wave dynamics in excitable media, appears to be unexpectedly non-obvious and understudied (see below). In this paper, we prove the aforementioned consequence using the remarkable features of optogenetics. Our study is one of the first to demonstrate the establishment of real-time spatiotemporal control over spiral wave dynamics in spatially extended biological (cardiac) tissue. The only other study that shows effective dynamic spatiotemporal control (as opposed to elimination) of spiral waves is that of (***Burton et al., 2015***). However, (***Burton et al., 2015***) use dynamic control to modulate spiral wave chirality, which is markedly different from what we study and prove, namely spiral wave dragging.

In this research, we demonstrate a new phenomenological concept to steer spiral waves in excitable media, thereby enabling precise and direct spatiotemporal control over spiral wave dynamics. This so-called AAD control involves a feedback interaction between 3 fundamental dynamical properties of spiral waves: attraction, anchoring and unpinning. In the past, researchers also employed different kinds of spatiotemporal feedback interactions to control spiral wave dynamics, albeit in non-biological media (***Sakurai et al., 2002***; ***Schlesner et al., 2008***). Their methods, while effective, relied on delivering timed pulses falling in defined phases of the spiral. Although feasible in simple chemical systems, phase-based pulse delivery can be quite challenging in complex biological tissue, where temporal precision is a major subtlety and therefore difficult to achieve. The study by (***Ke et al., 2015***) bypasses this issue by forcing scroll wave filaments to anchor to physical heterogeneities, which can be moved subsequently to steer the waves. However, (***Ke et al., 2015***) demonstrate a process that is continuous in time, as opposed to the discrete nature of our method, which makes it more flexible.

Previous studies have explored the concept of spiral attraction in human cardiac tissue (***Defauw et al., 2014***). According to these findings, the mechanism of attraction of spiral waves to localized heterogeneities is a generic phenomenon, observed among spiral waves in heterogeneous tissue. Some studies (***Defauw et al., 2014***; ***Panfilov and Vasiev, 1991***; ***Rudenko and Panfilov, 1983***) show that spiral waves tend to drift to regions with longer rotational periods, i.e. where the duration of an action potential is longer than elsewhere in the medium. This is in consonance with our simulations, where application of a spot of light to a monolayer of excitable, optogenetically modified cardiomyocytes causes the cells in this region to depolarize, bearing a direct influence on the action potential duration of cells in the immediate neighborhood. Once attracted, the spiral anchors to the location of the light spot, thereby making a transition from functional to anatomical reentry. Again, when the light at the current location is switched off and a new location is illuminated, in the next step of the drag sequence, the spiral reverts back to functional reentry, from its anatomical variant. If the new location is such that the spiral core overlaps with the basin of attraction of the optogenetically depolarized area, then it anchors to the new location and makes a transition back to anatomical reentry. Thus, AAD control is associated with repeated alternative transitions between functional and anatomical reentry. However, if the core of the spiral wave does not overlap with the basin of attraction of the next location in the drag sequence, or, if the drag sequence is executed too fast for the spiral to establish a stable core at subsequent new locations, then the spiral detaches, breaks up, or destabilizes without being successfully dragged.

A mechanism, somewhat along the same lines as AAD control, was demonstrated by Krinsky et al. (***Krinsky et al., 1995***), using topological considerations. When an existing vortex interacts with a circular wave emerging from a stimulus applied close to the vortex core, they fail to annihilate each other. However, the emergent wave quenches the vortex, resulting in its displacement by the order of half a wavelength. This displacement may occur in any direction, as demonstrated by Krinsky et al. In our system, we observe similar emergence of a new (circular) wave from the temporal heterogeneity. This new wave fails to annihilate the existing spiral wave, in line with Krinsky’s findings, but leads to the displacement of the core towards the temporal heterogeneity, indicating the existence of a factor of attraction, which determines the direction of drift.

In another study (***Guo et al., 2010***), control of spiral wave turbulence was investigated in a selfregulation system that spontaneously eliminated vortex cores, by trapping the latter with localized inhomogeneities. This study used computer simulations and a light-sensitive BZ reaction with a Doppler instability. Movement of a spiral wave along a line of predefined obstacles was also observed by Andreev et al. (***Andreev et al., 2005***). However, unlike Krinsky et al. and Guo et al. (***Guo et al., 2010***), their study uses inactive defects, which do not lead to quenching of the spiral wave. Further topological considerations with respect to movement of spiral waves can be found in the seminal work of Winfree et al. (***Winfree and Strogatz, 1984***). In our studies of spiral wave dragging with light spots of different sizes (Figure 3), a small-sized spot leads to dragging via a cycloidal trajectory, whereas, a large spot is associated with a linear trajectory. In our subsequent investigations, however, cycloidal trajectories were observed even for large spots. This is because, in the data presented in Figure 3, we investigate dragging at the fastest possible rate. The drag trajectories obtained in that study apply for minimal relocation time (*τ*_*light*_ = *τ*_*min*_, *τ*_*dark*_ = 0 ms). The larger the value of *d*, the bigger is the area of light-induced depolarization. Consequently, the higher is the chance for the tip of the spiral wave to fall within the basin of attraction of the illuminated area, causing it to remain anchored to the spot at all times. When the spot of light is moved with *τ*_*dark*_ = 0 ms and *τ*_*light*_ = *τ*_*min*_, the location of the new light spot forms a composite anchor point in combination with the location of the previous light spot; the effective depolarized region resembles an elongated ellipse, with elongation along the direction of movement of the light spot. This causes the spiral tip to trace a nearly linear path, along the boundary of the elongated ellipse. However, at slower dragging rates, i.e. longer *τ*_*ight*_, as used in the subsequent studies (Figure 3), the spiral wave gets ample time to execute one or more complete rotations around the region of light-induced depolarization, before being relocated to the next spot. Hence, the cycloidal trajectory at large *d*. The tendency for the spiral tip to move along a cycloidal trajectory at small *d* can be explained by the establishment of a very small basin of attraction, which results in an increased demand for *τ*_*min*_. Thus the spiral requires execution of at least one complete rotation around the region of light-induced depolarization before it can relocate to the next spot.

These findings are in consonance with the fundamental view of excitable media as dynamical systems. Based on their experimental, computational and theoretical studies, (***Zykov et al., 2010***) and (***Nakouzi et al., 2016***) reported the involvement of bistability and hysteresis in the transitions from anchoring to unpinning of a wave rotating around an anatomical hole. Bistability, or coexistence of two dynamical attractors in the system, indicates dependence of the outcome of the transition on initial conditions. In our case, the most important choice of initial conditions can be restricted to the phase of the freely rotating spiral wave or the rigidly anchored reentrant wave. Thus, proper timing of the applied perturbation is crucial regarding the efficacy of control. In this sense our results deceptively remind us of the phenomenon of spiral wave drift, which can be induced by periodic modulation of excitability (***Steinbock et al., 1993***). However, such drift arises from an interplay between Hopf bifurcation of spiral wave solutions, and rotational and translationally symmetric planar (Lie) groups (***Wulff, 1996***). In our studies, we rely on completely different nonlinear dynamical phenomena, namely, bistability of the rotating wave solutions and its interplay with spatial and temporal degrees of freedom of the dynamically applied light spot (i.e., its position, size, period and timing). This combination gives rise to novel interesting spiral wave dynamics, which opens broad avenues for further theoretical and experimental investigation.

Although apparently straightforward, and, to some extent, predictable on the basis of generic spiral wave theory, our results are not at all obvious. Compared to other systems, such as the BZ reaction, cardiac tissue (i) possesses an inherently irregular discrete cellular structure, underlying the absence of spatial symmetries, (ii) lacks inhibitor diffusion and (iii) has different dimensions. Moreover, since the probability of attraction of a spiral wave to a heterogeneity is determined by the initial conditions and by the strength of successive perturbations, it is not a foregone conclusion that discrete application of light spots outside the core of a spiral wave should always end up in an anchored state. By showing, in cultured cardiac tissue, that spiral wave dynamics can be controlled by targeting the spiral wave core, we have been able to experimentally confirm pre-existing ideas about the behavior of spiral waves in complex systems.

Previous studies also demonstrated alternative methods to ‘control’ spiral-wave dynamics in excitable media, for example, by periodic forcing to induce resonant drift (***Agladze et al., 1987***; ***Biktasheva et al., 1999***; ***Agladze et al., 2007***), or by periodic modulation of excitability to manipulate spiral tip trajectories (***Steinbock et al., 1993***). These studies do demonstrate the potential to drive a spiral wave out of an excitable medium, or to force spiral waves to execute complex meander patterns, thereby making landmark contributions to the knowledge of pattern formation or even to cardiac arrhythmia management. However, the principal limitation of these methods is that they are indirect, giving reasonable control over the initial and final states of the system, with little or no control in between. It is herein where lies the advantage of AAD control, which allows precise and complete spatiotemporal control over spiral wave dynamics in two-dimensional excitable media.

### Clinical translation

Since we demonstrate AAD control method in a cardiac tissue system, a logical question would be, how to envision the application of this principle to the real heart in order to treat arrhythmias? Currently this topic faces major challenges. The practical application of optogenetics in cardiology is, in itself, a debatable issue. However, with recent advances in cardiac optogenetics (***Nussinovitch and Gepstein, 2015***; ***Crocini et al., 2016***; ***Nyns et al., 2017***; ***Bruegmann et al., 2018***; ***Boyle et al., 2018***), the future holds much promise. Firstly, we envision the usage of AAD control method in treating arrhythmias that are associated with scroll waves. Since the penetration depth for 470 nm light in cardiac tissue is rather short (500*µ*m) (***Bruegmann et al., 2016***), full transmural illumination is challenging, particularly in ventricles of large mammals like pig, monkey or human. There, AAD control may provide a powerful tool to regulate scroll wave dynamics by epicardial or endocardial illumination. Furthermore, we expect the method to prove most useful when dealing with ‘hidden’ spiral waves, i.e., spiral waves in remote locations of the heart that are unaccessible by ablation catheters. Ideally, one should build upon the concept introduced by (***Entcheva and Bub, 2016***). With live spacetime optogenetic actuation of the electrical activity in different parts of the heart, the first step is to detect the location of the instability. Next, one can use a catheter with an in-built LED to attract the scroll wave filament and steer it towards the nearest tissue border for termination. The advantage of this method lies in that one does not require to ablate, and thereby destroy, excitable cardiac tissue, thus avoiding the possibility to create permanent damage to the heart that may favor development and of new arrhythmias. In addition, as our study demonstrates, anchoring of the spiral core (scroll filament) can occur even if the ‘precise’ location of the core is not identified. Lastly, the discrete nature of our method allows temporal flexibility in steering the spiral core (scroll filament), in that, temporary loss of communication between the catheter and the spiral core will not lead to failure of the technique in general. The reversible nature of the AAD control technique makes it unsuitable for terminating arrhythmias that rely on the establishment of a permanent conduction block as can be produced via conventional catheter ablation. However, this special feature of AAD may come with certain unique advantages that should be explored in more detail in future studies. For example, in younger patients that are expected to undergo periodic repetitive ablation for termination of reoccurring arrhythmias of unknown origin, the non-destructive nature of AAD may prove to be more desirable than the cumulative widespread destruction of cardiac tissue by radiofrequency or cryoballoon ablation. Alternatively, other methods could be developed and explored for AAD control without the need of optogenetic modification, while still relying on the creation of spatiotemporally controlled heterogeneities for attraction, anchoring and dragging of spiral waves.

In this study, we focus on spiral waves in cardiac excitable media, as these abnormal waves have been associated with lethal heart rhythm disturbances, while their management and termination remain a serious challenge. The insights gained from our results, as well as the AAD control method itself, may not only improve our understanding of spiral waves dynamics in favor of restoring normal cardiac rhythm, but also create incentive to explore these principles in other excitable media prone to spiral wave development.

## Methods and Materials

The electrophysiological properties of neonatal rat atrial cardiomyocytes was modeled according to (***Majumder et al., 2016***), whereby, the transmembrane potential ***V*** of a single cell evolved in time as follows:

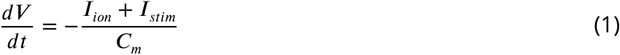

*I*_*ian*_ is the total ionic current, expressed as a sum of 10 major ionic currents, namely, the fast *Na*^+^ current (*I*_*Na*_), the ***L* - *type Ca*^2+^** current (*I*_*CaL*_), the inward rectifier *K*^+^ current (*IK*1), the transient outward *K*^+^ current (_*Ita*)_, the sustained outward *K*^+^ current (*I*_*Ksus*_), the background *K*^+^ (*I*_*Kb*_), the background *Na*^+^ (*I*_*Nab*_) and the background *Ca*^2+^ (*I*_*Cab*_) currents, the hyperpolarization-activated funny current (*If*), and the acetylcholine-mediated *K*^+^ current (*I*_*K,ACh*_). In two dimensions, these cells were coupled such that *V* evolved spatiotemporally, obeying the following reaction-diffusion equation:

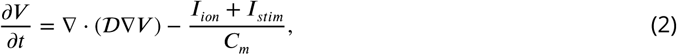

where 𝒟 is the diffusion tensor. In simple two-dimensional (2D) monolayer systems, in the absence of anisotropy, the diffusion tensor takes on a diagonal form with identical elements. Thus 𝒟 reduced to a scalar in our calculations, with value 0.00012 cm^2^/ms. This resulted in a signal conduction velocity of 22.2 ± 3.4 cm/s. Furthermore, in 2D, *I*_*K,ACh*_ was considered to be constitutively active, in consonance with the results from *in vitro* experiments (***Majumder et al., 2016***).

The optogenetic tool used in the numerical studies was a model of *Chlamydomonas reinhardtii* channelrhodopsin-2 mutant H134R, adopted from the studies of (***Boyle et al., 2013***) The parameter set used in our studies was exactly the same as that reported by (***Boyle et al., 2013***), with irradiation intensity 3.0 mW/mm^2^ to qualitatively mimic the large photocurrent produced upon illuminating a neonatal rat atrial cardiomyocyte monolayer.

In order to consistency with the *in vitro* experiments, we prepared 10 different simulation domains, composed of neonatal rat atrial cardiomyocytes with 17% randomly distributed cardiac fibroblasts (***MacCannell et al., 2007***) and with natural cellular heterogeneity, modelled as per (***Majumder et al 2016***). Our analysis was performed by taking into consideration, the results obtained from all 10 *in silico* monolayers. In order to demonstrate phenomena such as manipulation of spiral core size and position, we designed several ‘illumination’ patterns, in the form of a sequence of filled circles of desired radii and projected these patterns on to each of our *in silico* monolayers. Depending on the aim of the study, we adjusted the durations for which the light was ‘on’ (i.e. light interval), or ‘off’ (i.e. dark interval).

### Preparation of CatCh-expressing monolayers

All animal experiments were reviewed and approved by the Animal Experiments Committee of the Leiden University Medical Center and conformed to the Guide for the Care and Use of Laboratory Animals as stated by the US National Institutes of Health. Monolayers of neonatal rat atrial cardiomyocytes expressing Ca^2+^-translocating channelrhodopsin (CatCh) were established as previously described (***Feola et al., 2017***). Briefly, the hearts were excised from anesthetized 2-day-old Wistar rats. The atria were cut into small pieces and dissociated in a solution containing 450 U/ml collagenase type I (Worthington, Lakewood, NJ) and 18,75 Kunitz/ml DNase I (Sigma-Aldrich, St. Louis, MO). The resulting cell suspension was enriched for cardiomyocytes by preplating for 120minutes in a humidified incubator at 37^°^C and 5% CO_2_ using Primaria culture dishes (Becton Dickinson, Breda, the Netherlands). Finally, the cells were seeded on round glass coverslips (*d* = 15 mm;Gerhard Menzel, Braunschweig, Germany) coated with fibronectin (100 *µ*g/ml; Sigma-Aldrich) to establish confluent monolayers. After incubation overnight in an atmosphere of humidified 95%air5% CO at 37^°^C, these monolayers were treated with Mitomycin-C (10 *µ*g/ml; Sigma-Aldrich)for 2 hours to minimize proliferation of the non-cardiomyocytes. At day 4 of culture, the neonatal rat atrial cardiomyocyte monolayers were incubated for 20-24 h with CatCh-encoding lentiviral particles at a dose resulting in homogeneous transduction of nearly 100% of the cells. Next, the cultures were washed once with phosphate-buffer saline, given fresh culture medium and kept under culture conditions for 3-4 additional days.

### Optical mapping and optogenetic manipulation

Optical voltage mapping was used to investigate optogenetic manipulation of spiral wave dynamics in the CatCh-expressing monolayers on day 7 of culture by using the voltage-sensitive dye di-4-ANBDQBS (52.5 *µ*M final concentration; ITK diagnostics, Uithoorn, the Netherlands), as described previously (***Feola et al., 2017***). In summary, optical data were acquired using a MiCAM ULTIMA-L imaging system (SciMedia, Costa Mesa, CA) and analyzed with BrainVision Analyzer 1101 software (Brainvision, Tokyo, Japan). Only monolayers characterized by uniform AP propagation at 1-Hz pacing and homogeneous transgene expression were included for the following optogenetic investigation. CatCh was locally activated by using a patterned illumination device (Polygon400; Mightex Systems, Toronto, ON) connected to a 470-nm, high-power collimator light-emitting diode (LED) source (50 W, type-H, also from Mightex Systems). PolyLite software (Mightex Systems) was used to control the location and movement of the areas of illumination. Before local optogenetic manipulation, reentry was induced by light-based stimulations (n=18). Reentrant waves that were stable for >1 s were exposed to a circular light spot (*d* = 3 mm) at different targeted locations within the monolayer for 200 - 250 ms at 0.3 mW/mm^2^.

## Supplemental figure legends

AAD control of spiral wave cores to accomplish termination of a complex figure-of-eight type reentry, where the constituting spirals rotate slightly out of phase with each other. (A1) Positions of phase singularities of the initial reentrant pattern are marked with red asterisks. (A2) Phase correction of spirals by anchoring to light spots of different sizes. (A3-5) Formation of a single multi-arm spiral. (A6-9) Dragging the multi-arm spiral to termination. In each frame, the current position of the light spot is marked with a filled blue circle. The approximate trajectory of the dragged reentrant pattern is shown with dashed red lines.

## Video legends

**Image processing for the videos *in vitro A*:** Optical voltage signal was processed with spatial and temporal derivative filters to allow visualization of electrical wave propagation during the application of the light spots.

**Image processing for the videos *in vitro B*:** The data were sampled according to the location of the applied light spot. Next, for each sample set, we obtained the lowest background intensity recorded by the individual pixels over time and constructed a two-dimensional array with these values as elements. Next, we subtracted this array from each frame in the sample set and reassembled the processed datasets to make a complete video.

**Video 1:** Use of AAD control method to drag a spiral wave core along a triangular trajectory in silico and in vitro. ***In vitro A*** emphasizes the shape of the wavefront. ***In vitro B*** shows the same data, but processed to show more clearly, what happens during the application of the light spots.

**Video 2:** Use of AAD control method to drag a spiral wave core to termination *in silico* and *in vitro*. ***In vitro A*** emphasizes the shape of the wavefront. ***In vitro B*** shows the same data, but processed to show more clearly, what happens during the application of the light spots.

**Video 3**: Use of AAD control method to terminate figure-of-eight type of reentry *in silico* and *in vitro*. ***In vitro A*** emphasizes the shape of the wavefront. ***In vitro B*** shows the same data, but processed to show more clearly, what happens during the application of the light spot.

**Video 4**: Use of AAD control method to terminate complex reentry *in silico*, with 3 (left) and 4(right) spiral waves.

**Video 5**: Use of AAD control method to terminate complex reentry (4 spiral waves) *in vitro*. This video was produced in the same manner as described above for ***In vitro A***.

## Acknowledgments

This study was supported by The Netherlands Organisation for Scientific Research (Vidi grant 91714336) and the European Research Council (Starting grant 716509), both to DAP. Additional support was provided by Ammodo (to DAP and AAFdV). RM would like to thank Dr. Patrick Boyle for discussions. IF would like to thank Cindy Bart and Annemarie Kip for their assistance with the animal experiments and lentiviral particles production.

---

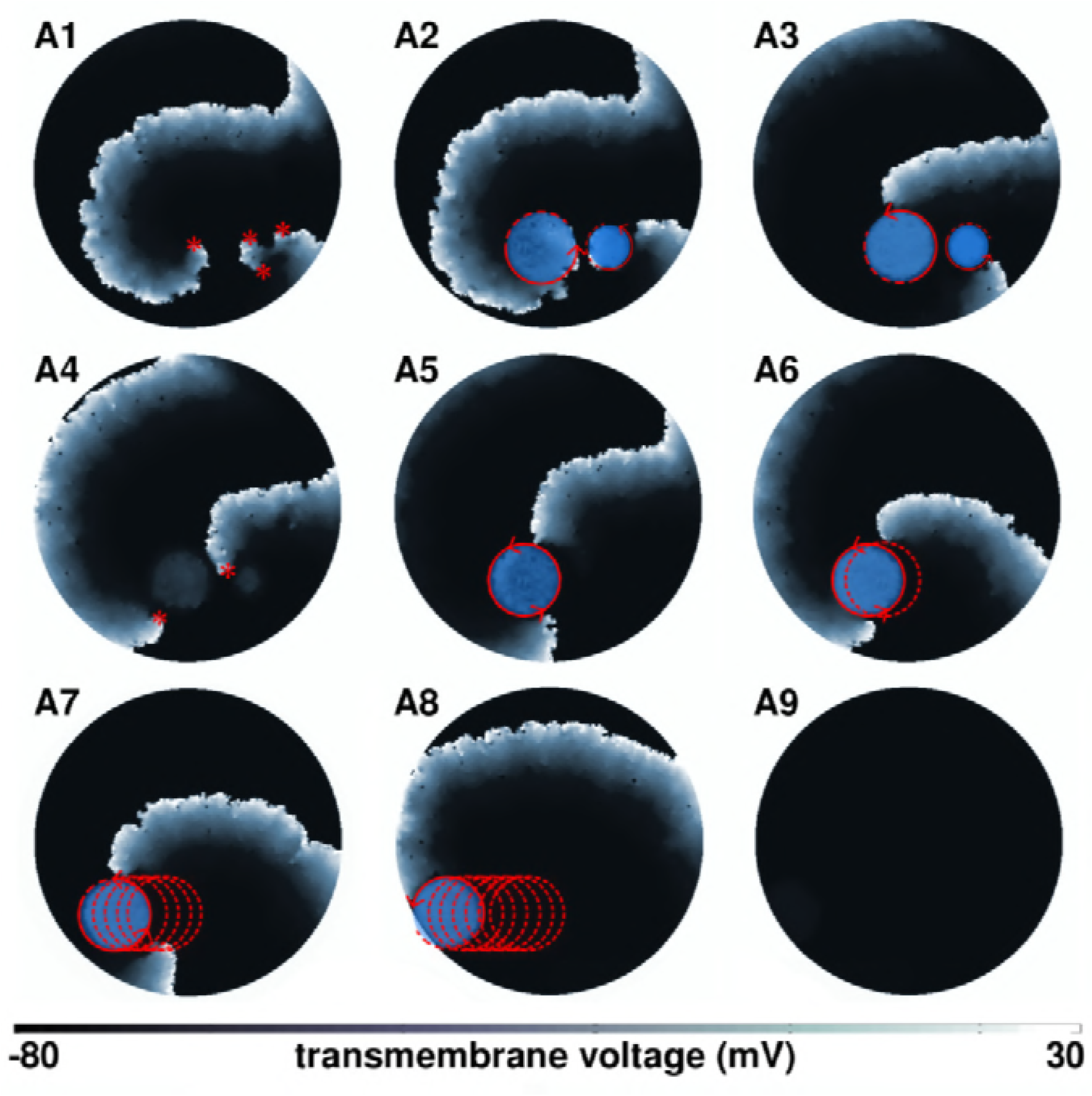

## References

Agladze KI, Davydov VA, Mikhailov AS. Observation of a helical wave resonance in an excitable distributed medium. JETP Lett. 1987; 45:767–770.

Agladze K, Kay MW, Krinsky V, Sarvazyan N. Interaction between spiral and paced waves in cardiac tissue. American Journal of Physiology-Heart and Circulatory Physiology. 2007; 293(1):H503–H513. doi:10.1152/ajpheart.01060.2006.

Andreev KV, Krasichkov LV. On the Possibility of Controlling the Motion of Spiral Waves in a Two-Dimensional Lattice of Excitable Elements with the Aid of Topological Defects. Technical Physics Letters. 2005; 31(7):545–47. doi: https://doi.org/10.1134/1.2001049.

Arrenberg AB, Stainier DY, Baier H, Huisken J. Optogenetic control of cardiac function. Science. 2010; 330(6006):971–4. doi:doi:10.1126/science.1195929.

Beaumont J, Davidenko N, Davidenko JM, Jalife J. Spiral Waves in Two-Dimensional Models of Ventricular Muscle: Formation of a Stationary Core. Biophysical Journal. 1998; 75:1–14. doi:doi:10.1016/S0006-3495(98)77490-9.

Belousov BP. In: A periodic reaction and its mechanism Wiley, New York; 1985.

Bi A, Cui J, Ma YP, Olshevskaya E, Pu M, Dizhoor AM, Pan ZH. Ectopic expression of a microbial-type rhodopsin restores visual responses in mice with photoreceptor degeneration. Neuron. 2006; 50(1):23–33.

Biktasheva IV, Elkin YE, Biktashev VN. Resonant Drift of Spiral Waves in the Complex Ginzburg-Landau Equation. J Biol Phys. 1999; 25(2-3):115–127. doi:10.1023/A:1005134901624.

Bingen BO, Engels MC, Schalij MJ, Jangsangthong W, Neshati Z, Feola I, Ypey DL, Askar SFA, Panfilov AV, Pijnappels DA, de Vries AAF. Light-induced termination of spiral wave arrhythmias by optogenetic engineering of atrial cardiomyocytes. Cardiovascular Research. 2014; 104(1):194–205. http://dx.doi.org/10.1093/cvr/cvu179, doi:10.1093/cvr/cvu179.

Boyden ES, Zhang F, Bamberg E, Nagel DKG. Millisecond-timescale, genetically targeted optical control of neural activity. Nat Neurosci. 2005; 8(9):1263–8.

Boyle PM, Williams JC, Ambrosi CM, Entcheva E, Trayanova NA. A comprehensive multiscale framework for simulating optogenetics in the heart. Nat Commun. 2013; 4(2370). doi:doi:10.1038/ncomms3370.

Boyle PM, Karathanos TV, Trayanova NA. Cardiac Optogenetics: 2018. JACC: Clinical Electrophysiology. 2018; http://electrophysiology.onlinejacc.org/content/early/2018/01/22/j.jacep.2017.12.006, doi:10.1016/j.jacep.2017.12.006.

Bruegmann T, Beiert T, Vogt CC, Schrickel JW, Sasse P. Optogenetic termination of atrial fibrillation in mice. Cardiovasc Res. 2018; 114(5):713–723. doi:doi:10.1093/cvr/cvx250.

Bruegmann T, Malan D, Hesse M, Beiert T, Fuegemann CJ, Fleischmann BK, Sasse P. Optogenetic control of heart muscle in vitro and in vivo. Nat Methods. 2010; 7(11):897–900. doi:doi:10.1038/nmeth.1512.

Bruegmann T, Boyle PM, Vogt CC, Karathanos TV, Arevalo HJ, Fleischmann BK, Trayanova NA, Sasse P. Optogenetic defibrillation terminates ventricular arrhythmia in mouse hearts and human simulations. The Journal of Clinical Investigation. 2016 10; 126(10):3894–3904. https://www.jci.org/articles/view/88950, doi:10.1172/JCI88950.

Burton RAB, Klimas A, Ambrosi CM, Tomek J, Corbett A, Entcheva E, Bub G. Optical control of excitation waves in cardiac tissue. 2015; 9:813–816. http://dx.doi.org/10.1038/nphoton.2015.196, doi:10.1038/nphoton.2015.196.

Caspi Y, Dekker C. Mapping out Min protein patterns in fully confined fluidic chambers. eLife. 2016; 5(e19271). doi:10.7554/eLife.19271.

Crocini C, Ferrantini C, Coppini R, Scardigli M, Yan P, Loew LM, Smith G, Cerbai E, Poggesi C, Pavone FS, Sacconi L. Optogenetics design of mechanistically-based stimulation patterns for cardiac defibrillation. Sci Rep. 2016; 6:35628. doi:doi:10.1038/srep35628.

Cross MC, Hohenberg PC. Pattern formation outside of equilibrium. Rev Mod Phys. 1993 jul; 65:851–1112. https://link.aps.org/doi/10.1103/RevModPhys.65.851, doi:10.1103/RevModPhys.65.851.

Davidenko JM, Kent P, Chialvo DR, Michaels DC, Jalife J. Sustained vortex-like waves in normal isolated ventricular muscle. Proc Natl Acad Sci USA. 1990; 87:8785–8789. doi: https://doi.org/10.1073/pnas.87.22.8785.

Davidenko JM, Kent P, Jalife J. Spiral waves in normal isolated ventricular muscle. Physica D. 1991; 49:182–197. doi: https://doi.org/10.1016/0167-2789(91)90207-P.

Defauw A, Vandersickel N, Dawyndt P, Panfilov AV. Small size ionic heterogeneities in the human heart can attract rotors. American Journal of Physiology-Heart and Circulatory Physiology. 2014; 307(10):H1456–H1468. doi:10.1152/ajpheart.00410.2014.

Deisseroth K. Optogenetics: 10 years of microbial opsins in neuroscience. Nat Neurosci. 2015; 18(9):1213–25. doi:doi:10.1038/nn.4091.

Entcheva E, Bub G. All-optical control of cardiac excitation: combined high-resolution optogenetic actuation and optical mapping. The Journal of Physiology. 2016; 594(9):2503–2510. https://physoc.onlinelibrary.wiley.com/doi/abs/10.1113/JP271559, doi:10.1113/JP271559.

Ermakova EA, Pertsov AM, Shnol EE. On the interaction of vortices in two-dimensional active media. Physica D: Nonlinear Phenomena. 1989; 40(2):185–195. http://www.sciencedirect.com/science/article/pii/0167278989900626, doi: https://doi.org/10.1016/0167-2789(89)90062-6.

Feola I, Volkers L, Majumder R, Teplenin A, Schalij MJ, Panfilov AV, de Vries AAF, Pijnappels DA. Localized Optogenetic Targeting of Rotors in Atrial Cardiomyocyte Monolayers. Circ Arrhythm Electrophysiol. 2017; 10:e005591. doi:10.1161/CIRCEP.117.005591.

Guo W, Qiao C, Zhang Z, Ouyang Q,Wang H. Spontaneous suppression of spiral turbulence based on feedback strategy. Phys Rev E. 2010; 81(056214). doi:10.1103/PhysRevE.81.056214.

Jia Z, Valiunas V, Lu Z, Bien H, Liu H, Wang HZ, Rosati B, Brink PR, Cohen IS, Entcheva E. Stimulating cardiac muscle by light: cardiac optogenetics by cell delivery. Circ Arrhythm Electrophysiol. 2011; 4(5):753–60. doi:doi:10.1161/CIRCEP.111.964247.

Karman TV. The Fundamentals of the Statistical Theory of Turbulence. Journal of the Aeronautical Sciences. 1937; 4:131–138. doi: https://doi.org/10.2514/8.350.

Kastberger G, Schmelzer E, Kranner I. Social Waves in Giant Honeybees Repel Hornets. PLOS ONE. 2008 09; 3(9):1–16. https://doi.org/10.1371/journal.pone.0003141, doi:10.1371/journal.pone.0003141.

Ke H, Zhang Z, Steinbock O. Scroll waves pinned to moving heterogeneities. Phys Rev E. 2015 Mar; 91:032930. https://link.aps.org/doi/10.1103/PhysRevE.91.032930, doi:10.1103/PhysRevE.91.032930.

Krinsky V, Plaza F, Voignier V. Quenching a rotating vortex in an excitable medium. Physical Review E. 1995; 52(3):2458–2462. doi:10.1103/PhysRevE.52.2458.

Krinsky VI. Mathematical models of cardiac arrhythmias (spiral waves). Pharmacol Ther B. 1978; 3:539–555. doi: https://doi.org/10.1016/S0306-039X(78)90020-X.

MacCannell KA, Bazzazi H, Chilton L, Shibukawa Y, Clark RB, Giles WR. A mathematical model of electrotonic interactions between ventricular myocytes and fibroblasts. Biophys J. 2007; 92(11):4121–32. doi:10.1529/biophysj.106.101410.

Majumder R, Jangsangthong W, Feola I, Ypey DL, Pijnappels DA, Panfilov AV. AMathematicalModel of Neonatal Rat Atrial Monolayers with Constitutively Active Acetylcholine-Mediated *K*^+^ Current. PLoS Comp Biol. 2016; 12(6):e1004946. doi: https://doi.org/10.1371/journal.pcbi.1004946.

McNamara HM, Zhang H, Werley CA, Cohen AE. Optically Controlled Oscillators in an Engineered Bioelectric Tissue. Phys Rev X. 2016 Jul; 6:031001. https://link.aps.org/doi/10.1103/PhysRevX.6.031001, doi:10.1103/PhysRevX.6.031001.

Mikhailov AS, Loskutov AY. Foundations of synergetics II: Chaos and Noise, vol. 52. Springer Science & Business Media; 2013.

Mikhailov AS, Showalter K. Control of waves, patterns and turbulence in chemical systems. Physics Reports. 2006; 425(2):79–194. http://www.sciencedirect.com/science/article/pii/S0370157305004825, doi: https://doi.org/10.1016/j.physrep.2005.11.003

Nakouzi E, Totz JF, Zhang Z, Steinbock O, Engel H. Hysteresis and drift of spiral waves near heterogeneities: From chemical experiments to cardiac simulations. Phys Rev E. 2016 Feb; 93:022203. https://link.aps.org/doi/10.1103/PhysRevE.93.022203, doi:10.1103/PhysRevE.93.022203.

Nussinovitch U, Gepstein L. Optogenetics for in vivo cardiac pacing and resynchronization therapies. Nature Biotechnology. 2015; 33:750–754. http://dx.doi.org/10.1038/nbt.3268, doi:10.1038/nbt.3268.

Nyns ECA, Kip A, Bart CI, Plomp JJ, Zeppenfeld K, Schalij M, De Vries A, Pijnappels DA. Optogenetic termination of ventricular arrhythmias in the whole heart: towards biological cardiac rhythm management. Eur Heart J. 2017; (27):2132–2136. doi:doi:10.1093/eurheartj/ehw574.

Panfilov AV, Vasiev BN. Vortex initiation in a heterogeneous excitable medium. Physica D. 1991; 49:107–113. doi:https://doi.org/10.1016/0167-2789(91)90200-S.

Ripplinger CM, Krinsky VI, Nikolski VP, Efimov IR. Mechanisms of unpinning and termination of ventricular tachycardia. Am J Physiol Heart Circ Physiol. 2006; 291:H184–H192. doi:10.1152/ajpheart.01300.2005.

Rudenko AN, Panfilov AV. Drift and interaction of vortices in two-dimensional heterogeneous active medium. Studia Biophysica. 1983; 98:183–188.

Sakurai T, Mihaliuk E, Chirila F, Showalter K. Design and Control of Wave Propagation Patterns in Excitable Media. Science. 2002; 296(5575):2009–2012. http://science.sciencemag.org/content/296/5575/2009, doi:10.1126/science.1071265.

Schebesch I, Engel H. Interacting spiral waves in the Oregonator model of the light-sensitive Belousov-Zhabotinskii reaction. Phys Rev E. 1999 Dec; 60:6429–6434. https://link.aps.org/doi/10.1103/PhysRevE.60.6429, doi:10.1103/PhysRevE.60.6429.

Schlesner J, Zykov VS, Brandtstädter H, Gerdes I, Engel H. Efficient control of spiral wave location in an excitable medium with localized heterogeneities. New Journal of Physics. 2008; 10(1):015003. http://stacks.iop.org/1367-2630/10/i=1/a=015003.

Schulman LS, Seiden PE. Percolation and Galaxies. Science. 1986; 233(4762):425–431. http://science.sciencemag.org/content/233/4762/425, doi:10.1126/science.233.4762.425.

Steinbock O, Müller SC. Light-controlled anchoring of meandering spiral waves. Phys Rev E. 1993 Mar; 47:1506–1509. https://link.aps.org/doi/10.1103/PhysRevE.47.1506, doi:10.1103/PhysRevE.47.1506.

Steinbock O, Schütze J, Müller SC. Electric-field-induced drift and deformation of spiral waves in an excitable medium. Phys Rev Lett. 1992 Jan; 68:248–251. https://link.aps.org/doi/10.1103/PhysRevLett.68.248, doi:10.1103/PhysRevLett.68.248.

Steinbock O, Zykov V, Müller S. Control of spiral-wave dynamics in active media by periodic modulation of excitability. Nature. 1993; 366:322–324. doi:doi:10.1038/366322a0.

Winfree AT, Strogatz SH. Organizing centres for three-dimensional chemical waves. Nature. 1984; 311(18):611–615. doi:doi:10.1038/311611a0.

Wulff C. Bifurcation theory of meandering spiral waves. In: Nonlinear Physics of Complex Systems Springer; 1996. p.166–178.

Zhabotinsky AM. A history of chemical oscillations and waves. Chaos. 1991; 1:379–86. doi:10.1063/1.165848.

Zykov V, Bordyugov G, Lentz H, Engel H. Hysteresis phenomenon in the dynamics of spiral waves rotating around a hole. Physica D: Nonlinear Phenomena. 2010; 239(11):797–807. http://www.sciencedirect.com/science/article/pii/S0167278909002383, doi:https://doi.org/10.1016/j.physd.2009.07.01, emergent Phenomena in Spatially Distributed Systems. 18

